# Excitatory-inhibitory windows shape coherent neuronal dynamics driven by optogenetic stimulation in the primate brain

**DOI:** 10.1101/437970

**Authors:** Ryan A. Shewcraft, Heather L. Dean, Margaret M. Fabiszak, Maureen A. Hagan, Yan T. Wong, Bijan Pesaran

## Abstract

Coherent neuronal dynamics play an important role in complex cognitive functions. Optogenetic stimulation promises to provide new ways to test the functional significance of coherent neural activity. However, the mechanisms by which optogenetic stimulation drives coherent dynamics remain unclear, especially in the non-human primate brain. Here, we perform computational modeling and experiments to study the mechanisms of optogenetic-stimulation-driven coherent neuronal dynamics in non-human primates. Neural responses arise from stimulation-evoked temporal windows of excitatory and inhibitory activity. The temporal properties of the E-I windows generate coherent neuronal dynamics at varied frequencies and depend on optogenetic stimulation parameters. Experimental results agree with parameter dependent predictions from the computational models. These results demonstrate that responses to optogenetic stimulation are governed by local circuit properties that alter the timing of E-I activity. Transient imbalances in excitatory and inhibitory activity may provide a general mechanism for generating coherent neuronal dynamics.

## Introduction

Spiking and local field potential (LFP) activity display coherent patterns of activity across a range of temporal frequencies (Pesaran et al., 2018). Coherent temporal structure has been proposed to play an important role in the control of higher cognitive functions (Buschman and Miller, 2007; Dean et al., 2012; Womelsdorf et al., 2006; Wong et al., 2016) and enhanced cortico-cortical communication more generally (Fries, 2005; Pesaran et al., 2018, 2008). Despite this, whether and how complex cognitive functions depend on the temporal structure of neuronal activity, as opposed to the magnitude of the activity, has been highly debated. Causal manipulations that generate frequency selective coherent neuronal dynamics are important for directly testing their functional significance. However, in order to interpret the results of causal manipulations we also need to understand how extrinsic modulations of coherent neuronal dynamics recruit intrinsic cellular and network mechanisms to generate the driven response.

In general, coherent neuronal dynamics depend on the temporal precision and synchrony of spike timing (Buzsáki and Wang, 2012), which is governed by the intrinsic dynamics of excitatory (E) and inhibitory (I) post-synaptic potentials (Brunel and Wang, 2003; Wehr and Zador, 2003). Typically, excitatory and inhibitory activity are tightly correlated, with E preceding I by several milliseconds (Okun and Lampl, 2008). Thus, inputs which drive excitatory activity are closely followed by inhibitory activity, forming a brief a window for temporally precise spiking in the postsynaptic neurons (Gabernet et al., 2005; Haider et al., 2013; Salkoff et al., 2015). The instantaneous frequency of coherent neuronal activity is regulated by E-I balance (Atallah and Scanziani, 2009) and depends on the relative timing of E and I activity (Chariker et al., 2018; Keeley et al., 2017). Consistent with this, empirically observed neural coherence is dynamic and episodic in the gamma (Burns et al., 2011, 2010; Xing et al., 2012) and beta (Feingold et al., 2015; Rule et al., 2017; Sherman et al., 2016) bands.

Optogenetic stimulation has been shown to alter neuronal dynamics extrinsically by manipulating either excitatory or inhibitory activity alone. In some studies, driving inhibitory activity by delivering pulse trains of optogenetic stimulation to interneurons perturbed coherent neuronal activity (Cardin et al., 2009; Sohal et al., 2009). In other studies, driving excitatory activity by stimulating pyramidal neurons using continuous optogenetic stimulation generated coherent dynamics, specifically gamma activity (Lu et al., 2015; Ni et al., 2016). However, coherent neuronal dynamics depend on the interaction between excitatory and inhibitory activity, which is governed by the intrinsic properties of the local circuit. Since stimulating either excitatory or inhibitory activity alone recruits both excitatory and inhibitory postsynaptic responses in the network (Buzsáki and Wang, 2012; Gloveli et al., 2005; Phillips and Hasenstaub, 2016; Pike et al., 2000; Tukker et al., 2007), optogenetic stimulation may generate coherent neuronal dynamics according to the relative timing of stimulation-driven E-I responses.

Motivated by this reasoning, we tested the hypothesis that optogenetic stimulation drives coherent neuronal dynamics by generating E-I responses that arise from intrinsic E-I balance. We first developed a computational model of coherent neuronal dynamics in response to optogenetic stimulation in terms of a temporal window of excitatory and inhibitory activity—the E-I window. We experimentally tested the model predictions by expressing ChR2(H134R) in macaque cortex delivered by an AAV2/5 viral injection and measuring spiking and local field potential (LFP) responses to sequences of optogenetic stimulation pulses. We demonstrate the generality of the findings across multiple regions of the posterior parietal and frontal cortex. Finally, we developed a spiking neural network model of the observed neural dynamics and found that E-I windows are an emergent property of the model. Together these results suggest that optogenetic stimulation extrinsically alters excitatory and inhibitory activity by recruiting intrinsic E-I windows to shape coherent neuronal dynamics in the beta and gamma frequency bands.

## Results

### Modeling optogenetic-stimulation-evoked coherent neuronal dynamics

Coherent neuronal dynamics arise from synchronous, temporally structured neuronal activity. Spike-field coherence (SFC) measures coherent neuronal dynamics by estimating the strength of correlations between spiking activity of individual neurons and neural population activity in the form of simultaneously-recorded LFP activity. When the spike times of a neuron can be predicted from fluctuations in LFP activity at a particular frequency, the spiking is said to be coherent with LFP activity at that frequency. Neurons that fire consistently at a particular phase of LFP activity are coherent with LFP activity. As a result, neuronal coherence is defined by coherent spiking and LFP activity and can be measured by estimating SFC between spiking and nearby LFP activity (Pesaran et al., 2018, 2002).

**Figure 1A** presents a model of optogenetic-stimulation-evoked coherent neuronal dynamics and corresponding idealized neural responses. During spontaneous activity, synaptic inputs drive activity in a local network. We model spontaneous spiking activity as being generated from a Poisson process with a constant rate (Brown et al., 2003) and spontaneous LFP activity as being generated from a Brown noise process. Under these assumptions, spontaneous spiking is not coherent with spontaneous LFP activity (**Figure 1A**, Spontaneous activity).

**Figure 1.**
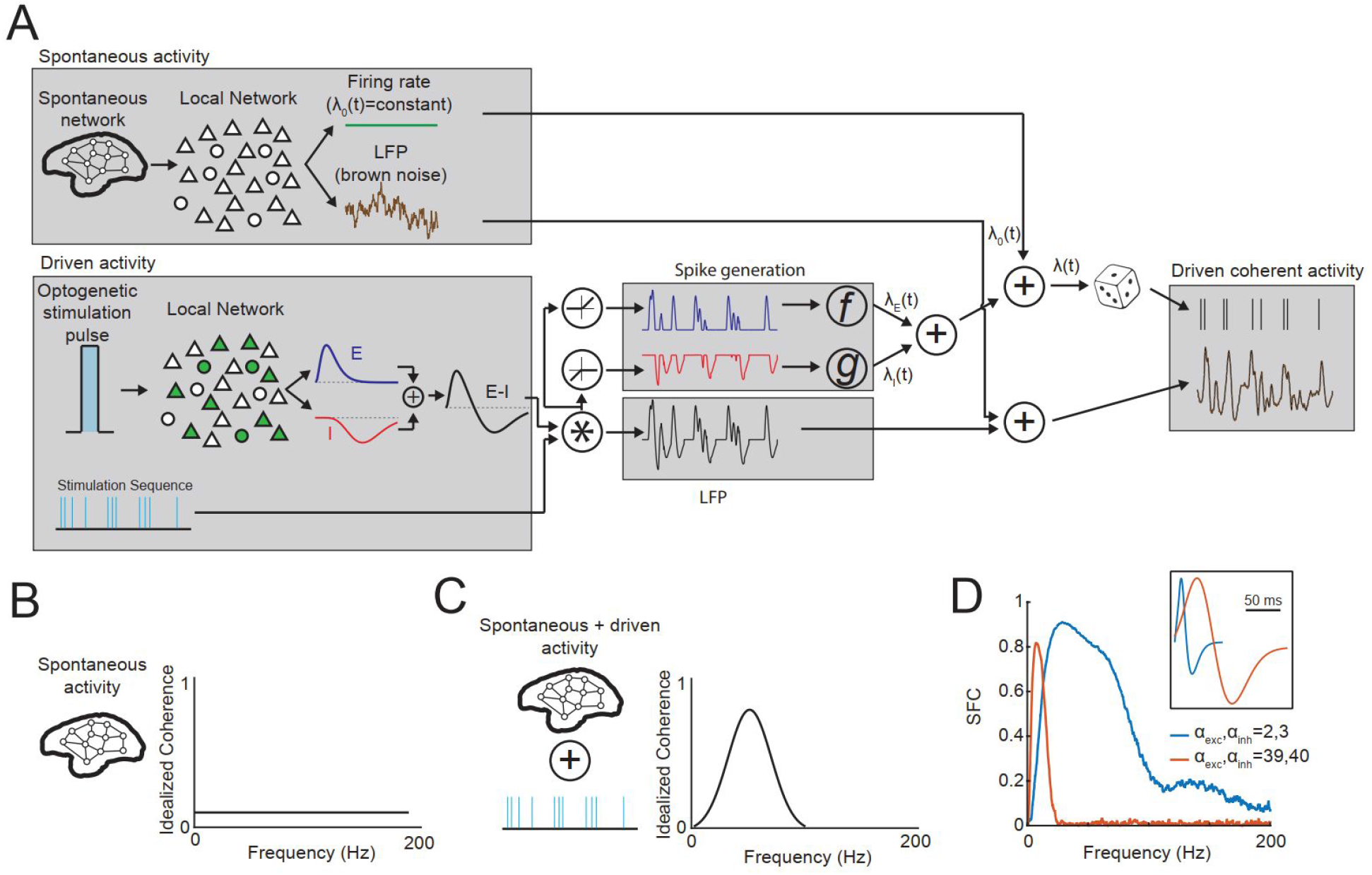
Excitation and inhibition drive coherent neuronal dynamics. **(A)** To model optogenetic stimulation of a population of transduced neurons, we simulated pulsatile responses generated by direct activation of ChR2 channels and synaptic activity. Pulsatile responses are convolved with a stimulation sequence. The summed excitatory and inhibitory (E-I, black) components comprise the simulated LFP. The individual E (blue) and I (red) activity govern the variable spiking rate in the Poisson process which generates coherent spiking. **(B)** Spontaneous activity in the model has no spike-field coherence. (C) Optogenetic stimulation drives correlated fields and spiking, leading to frequency selective spike-field coherence. **(D)** Simulated spike field coherence for short and long E-I windows. *Inset*, E-I windows used in the simulation.

Direct activation of transduced neurons leads to excitatory and inhibitory synaptic currents within the local network of neurons. The resulting excitatory (E) and inhibitory (I) currents in postsynaptic neurons occur at different moments in time and the relative timing of the E and I currents determines when neurons fire spikes (Cruikshank et al., 2010; Haider et al., 2013; Higley and Contreras, 2003; Wehr and Zador, 2003). Given that excitatory currents depolarize the membrane potential and inhibitory currents hyperpolarize the membrane potential, the cell has the greatest probability of firing when E is high and I is low. Therefore, we model the impact of optogenetic stimulation on spiking as an excitatory- and inhibitory-dependent temporal change in the probability of the cell firing (**Figure 1A**, Spike generation). Thus, the timing of spiking activity depends on the net postsynaptic E and I consequences of each stimulation pulse - the E-I window.

To simulate neuronal activity that exhibits SFC, we need to model the coupling between spiking and LFP activity. LFP activity predominantly reflects summed postsynaptic potentials (Buzsáki et al., 2012; Pesaran et al., 2018). Since the E and I postsynaptic activity drives spiking, we can model the driven LFP as the sum of the driven E and I responses, thus linking spiking and LFP activity,. We add brown noise background activity to the driven LFP activity to generate the simulated LFP (**Figure 1A**, LFP).

We refer to this model of neuronal coherence as the **E-I window model.** We model spontaneous spiking and LFP as uncorrelated so the spontaneous activity does not exhibit SFC (**Figure 1B**). Since optogenetic stimulation evokes E-I windows in which action potentials are more likely to occur, changes in spike probability are coupled to LFP responses. This coupling results in spikes that are coherent with LFP activity (**Figure 1C**). Importantly, the E-I window model predicts that the coherence is due to the temporal dynamics, or shape, of the E-I windows. In line with the prediction, SFC varied with the E-I window duration such that longer E-I windows generated SFC at lower frequencies (**Figure 1D**). In our empirically measured E-I windows, excitatory responses dominated the early stage followed by inhibitory responses. This is consistent with sensory stimulation which typically generates an increase in excitation followed shortly by a corresponding inhibitory response (Higley and Contreras, 2003; Wehr and Zador, 2003). However, inhibition may be stronger and have faster rise times than excitation in certain circuits or behaviors (Cruikshank et al., 2010; Haider et al., 2013). This regime would be captured in the E-I window model by setting *α_inh_* > *α_exc_*.

Note that the neuronal coherence described by the E-I window model depends on predictability between spikes and LFP activity. This construction is consistent with, but does not require, invoking an oscillator or other phase-consistent process. This construction is also consistent with the use of SFC, as opposed to other metrics of phase consistency, to estimate coherent neuronal dynamics. This is because SFC measures linear predictability between spikes and LFP activity. Note also that our approach contrasts with and is complementary to previous work that has used cell-type-specific optogenetic stimulation to generate neuronal coherence.

### Testing the E-I window model with optogenetic stimulation

Next, we tested whether optogenetic stimulation generates empirically observed neural coherence according to the predictions of the E-I window model. The model suggests that the frequency of driven coherent neuronal dynamics depends on the E-I window duration. Therefore, varying the duration of the stimulation pulse should change the E-I window duration, and the frequency content of the resulting driven coherent neuronal activity.

To test this hypothesis, we injected AAV5-hSyn-ChR2(H134R)-EYFP and recorded neuronal responses in three alert macaque monkeys quietly seated in a primate chair. We recorded in the inferior and superior parietal lobules in Monkeys H and J and lateral to the post-central dimple in Monkey B (**Figure 2A**). We observed strong viral expression at injection sites with robust labelling of neurons and neuropil (**Figure 2B**). We performed simultaneous *in vivo* optogenetic stimulation and recording using a fiber optic cable placed on a surgically thinned dura and adjacent to the intracortical recording electrode (**Figure 2C**). Stimulation consisted of one second long sequences of light pulses (stimulation epoch) followed by 1-3 s of no stimulation (spontaneous epoch) (**Figure 2D**). Optogenetic stimulation reliably generated neural responses. These responses consisted of action potentials as well as large evoked field potentials visible on an individual pulse basis in the raw data (**Figure 2E,F,G**).

**Figure 2.**
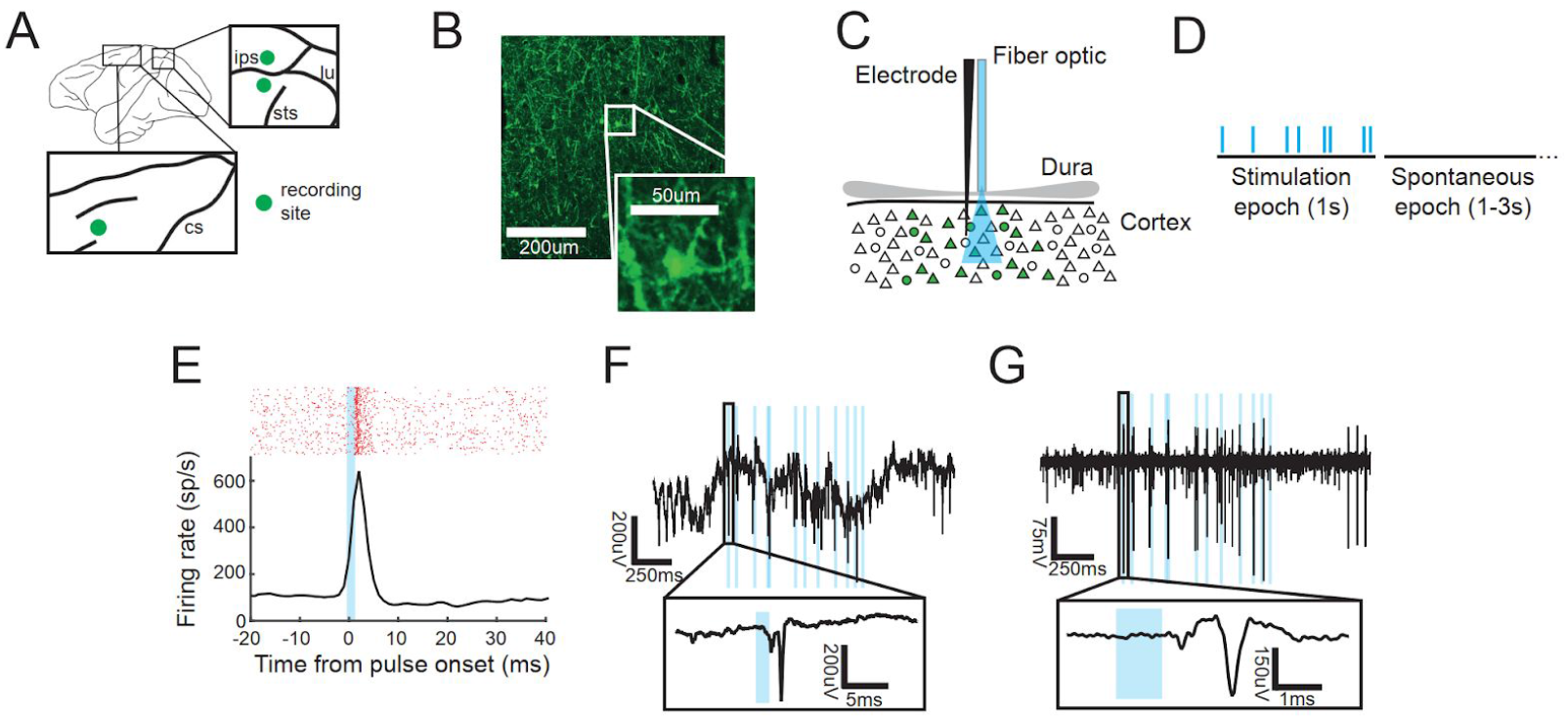
Optogenetic stimulation in macaque cortex. **(A)** Recording locations in all animals. **(B)** Example injection site with YFP labeled neurons and neuropil from a horizontal section. *Inset*. image of a single transduced neuron. (C) Recordings were performed with a single, acute electrode during trans-dural light stimulation with a fiber optic placed over a surgically thinned dura. **(D)** Experimental design featured a 1 s stimulation pulse sequence followed by 1-3 s of spontaneous activity without stimulation. **(E)** Example raster plot (top) and PSTH (bottom) of the spiking response triggered on each stimulation pulse for Poisson distributed trains of 1 ms pulses at 10 pulse/s rate. **(F)** Example broadband recording during a pulse train containing 1 ms pulses at 10 pulse/s rate; optogenetic stimulation (blue). *Inset*: Response to a single pulse. **(G)** Example multiunit filtered data from (E). *Inset*: Stimulation-elicited action potential.

We measured neuronal responses to stimulation sequences composed of Poisson pulse trains with different pulse rates and pulse durations. **Figure 3A** presents example neuronal responses during a stimulation block with 5 ms wide pulses delivered with a 20 pulse/s Poisson pulse train. Individual stimulation pulses drove spiking activity and evoked LFP responses at the stimulation site (**Figure 3Ai,ii**). We estimated the E-I windows by fitting the the E-I window to the empirically measured evoked LFP (**Figure 3Aiii**). We used this fitted E-I window in the E-I window model to simulate coherent neuronal dynamics. The simulated SFC closely matched the empirically measured SFC (**Figure 3Aiv**). This suggests that the E-I window model captures the phenomenology of the coherent neuronal dynamics driven by optogenetic stimulation. In addition, we computed the power spectrum of LFP activity during the one second long stimulation epoch and for one second of the spontaneous epoch (**Figure 3Av**). The stimulation LFP power spectrum revealed a significant increase in power (Chi-squared test, p<0.01).

**Figure 3.**
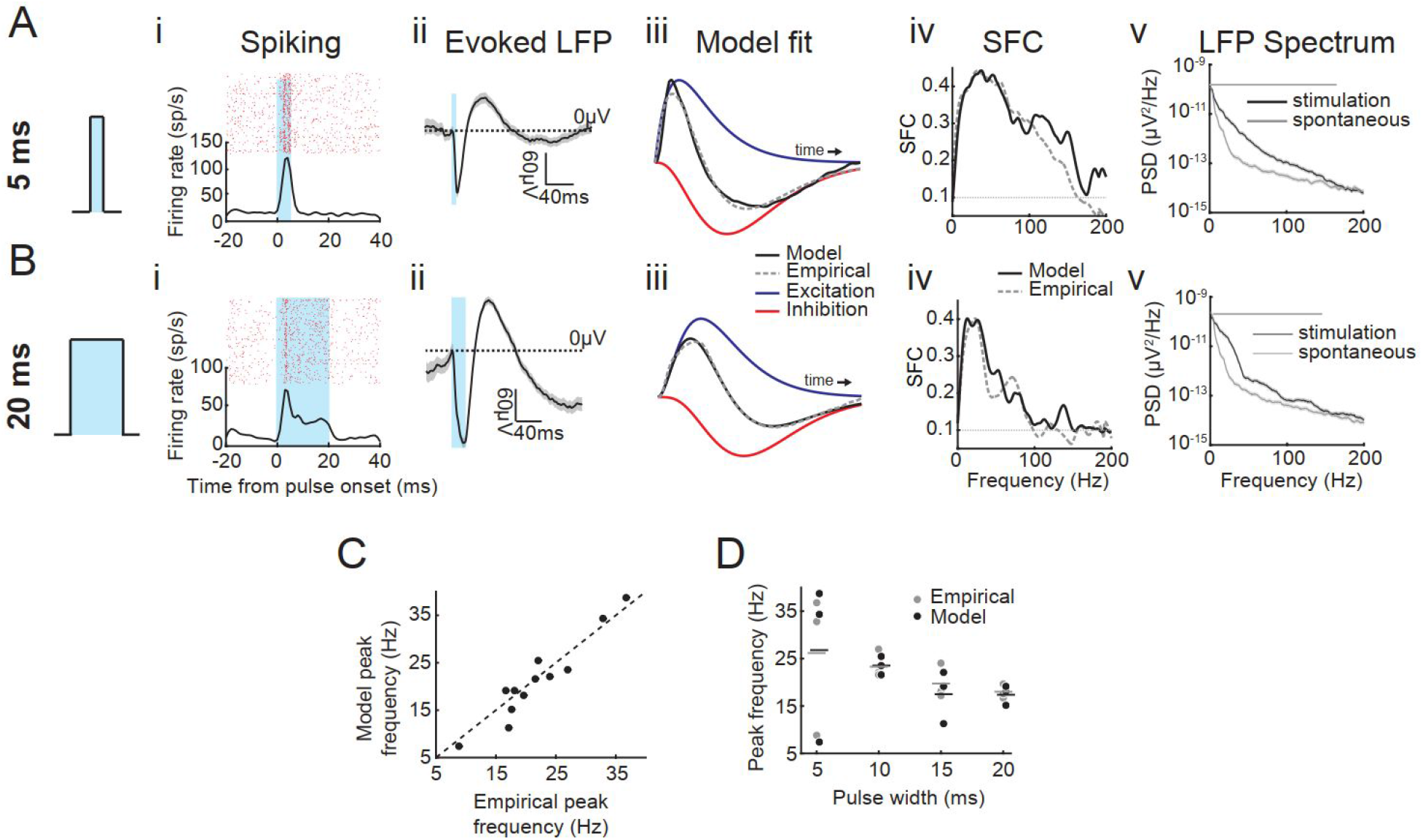
Neuronal dynamics driven by optogenetic stimulation. **(A)** Neuronal responses to 5 ms pulse duration, 20 Hz pulse rate, Possion pulse train. (i) Raster (top) and PSTH (bottom) of the response to single light pulses (blue). (ii) Mean evoked potential for all the stimulation pulses that were the last in a sequence and not preceded by a prior stimulation pulse within 50 ms, normalized by subtracting the value of the LFP at the onset of the light. Dotted line indicates 0 mV. (iii) Modelled E-I window (black) fit to the empirically measured E-I window (grey dotted) with excitatory (blue) and inhibitory (red) components. The fitted E-I window parameters are A=0.097, α_exc_=1, α_inh_=3, β_exc_=2.568, β_inh_=3.654.(iv) Spike-field coherence for multi-unit activity and LFP measured on the same electrode. Dotted horizontal line shows expected coherence under the null hypothesis that coherence is zero. (v) Power spectral density (PSD) of 1 s of data during the stimulation epoch (black line) and the spontaneous epoch (grey line). The dotted lines are error bars determined by the Chi-squared test for p=0.01. Solid gray lines above the PSD illustrate frequencies where the driven PSD was significantly different from the spontaneous PSD (p<0.01). **(B)** Same as (A) for 20 ms, 20 pulses/s pulse rate, Poisson pulse train. The fitted E-I window parameters for (Biii) are A=0.097, α_exc_=2, α_inh_=4, β_exc_=4.548, β_inh_=4.730. **(C)** Comparison of the peak frequency of the driven empirical and simulated SFC for all pulse widths conditions and all units. **(D)** Peak frequency for the driven SFC for each unit for the empirical data (grey) and E-I window model simulation (black). Horizontal lines show the mean for each condition.

Longer pulse durations generated longer duration changes in firing rate (**Figure 3Bi**) and longer periods of excitation and inhibition in the LFP activity (**Figure 3Bii,iii**), and have more selective changes in SFC and the LFP power spectrum (**Figure 3Bivv**). This shows that, stimulation sequences with different pulse durations can be used to selectively generate coherent neuronal dynamics, consistent with changing the E-I window duration. Thus, the E-I window model predicted the relationship between the E-I windows and the frequency content of the stimulation-evoked coherent neuronal dynamics.

### Optogenetic stimulation generates pulse duration-dependent responses

We computed the SFC for the population of recording sites with driven spiking activity in response to stimulation using Poisson pulse trains with 5, 10, 15, and 20 ms duration pulses. For each recording condition, we fit the E-I window as in Figures 3A and 3B and used the E-I window model to generate simulated SFC. For all recordings, the simulated SFC closely matched the empirical SFC (**Figure 3C**). In addition, for both the empirical and simulated data, the frequency of the driven SFC varied with pulse duration (**Figure 3D**), consistent with the example site and predictions from the E-I window model.

Importantly the center frequency did not depend on the mean pulse rate for Poisson stimulation (**Figure 3—figure supplement 1**). The decoupling of frequency from pulse rate means that the amount of coherence can be controlled separately from the center frequency. These results show that coherent neuronal dynamics in either beta or gamma frequency bands can be generated by stimulating the same population of neurons using stimulation pulses of different durations.

### Model-based investigation of excitatory and inhibitory dynamics

Since we did not have direct experimental access to excitatory and inhibitory currents, we developed a spiking neural network model to study how the frequency content of the driven coherent neuronal dynamics depended on the timing of the E and I activity. We implemented a computational model that consisted of a randomly-connected network of leaky integrate-and-fire excitatory and inhibitory neurons that captures LFP responses to sensory stimulation (Mazzoni et al., 2008). A subset of neurons also expressed ChR2 ion channels (Witt et al., 2013) (**Figure 4A,B**).

**Figure 4.**
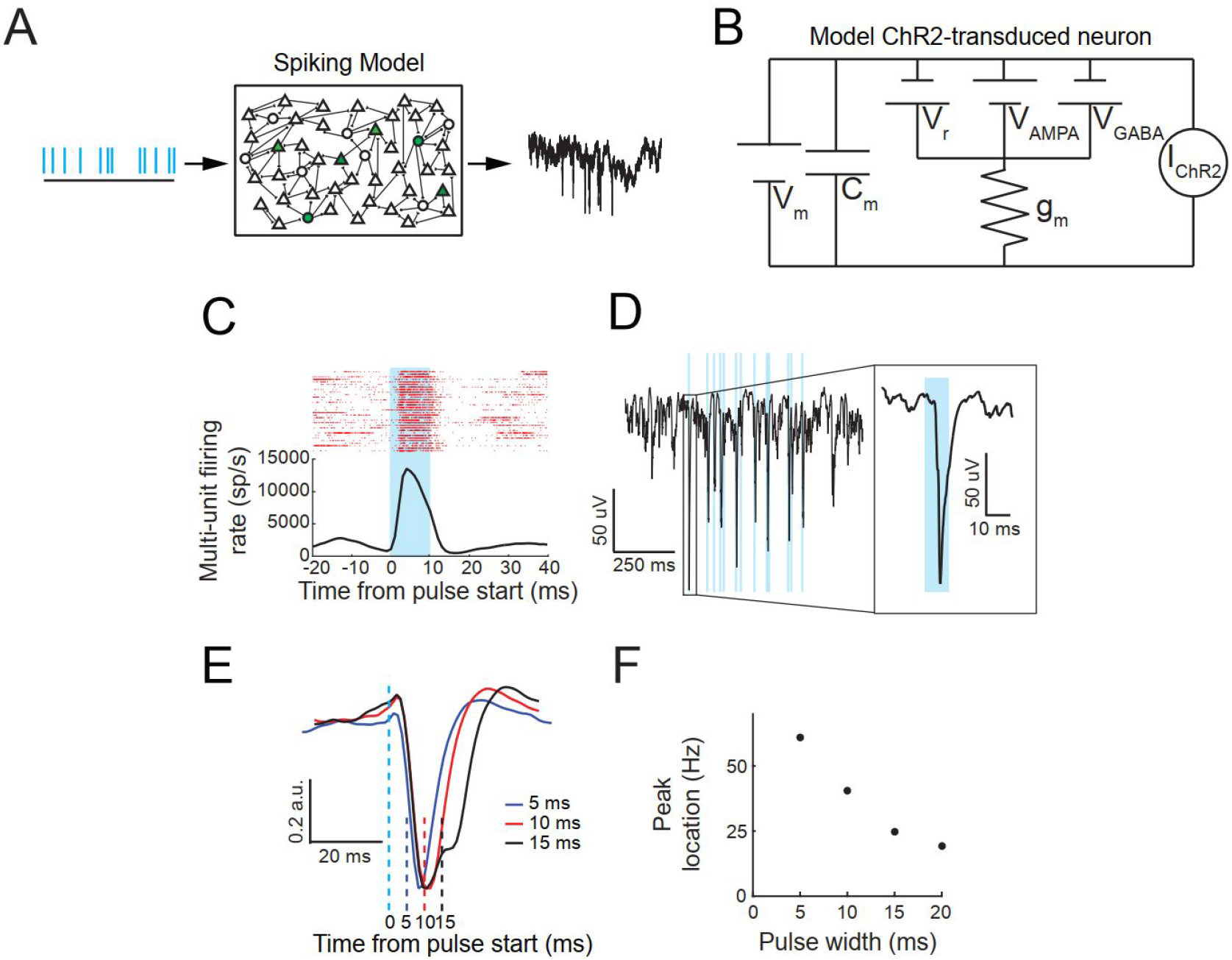
Spiking model responses and dynamics. **(A)** Diagram of a leaky integrate-and-fire model network with ChR2-transduced neurons. **(B)** Circuit diagram of leaky integrate-and-fire neurons with ChR2 channels that make up the spiking network model. **(C)** Raster (top) and PSTH (bottom) of the spiking model response to single light pulses for 10 ms, 10 pulses/s, Poisson stimulation. **(D)** Example broadband spiking model simulated LFP activity during a pulse sequence (10 ms, 10 pulses/s Poisson stimulation; blue lines indicate pulse times). *Inset*: Broadband simulated LFP activity at the time of a single pulse. **(E)** Mean normalized evoked potentials for the spiking model during pulse trains with 5 ms, 10 ms and 15 ms pulse durations and 10 pulses/s, Poisson stimulation. The dotted light blue line indicates the onset of the light stimulation and the subsequent dotted lines show the time when the light turned off. **(F)** Peak frequency of spiking model simulated SFC for each pulse width.

The model generated spiking activity and subthreshold potentials in response to optogenetic stimulation (**Figure 4C,D**). To test whether the model responses depended on stimulation pulse duration, we simulated data with the same pulse parameters as in **Figure 3** (5-20 ms pulse widths; 20 pulse/s Poisson). The model responses exhibit pulse duration dependence (**Figure 4E**) and frequency selectivity that varies with pulse duration (**Figure 4F**). Consistent with the empirical results, longer pulses durations resulted in a decrease in the center frequency of the driven coherent neuronal dynamics (**Figure 4F**). Therefore, the spiking model captures the key features of the empirically observed optogenetically-driven coherent neuronal dynamics. As with the empirical data and E-I window model, the frequency selectivity of the driven SFC is pulse-duration-dependent.

The spiking model responses depended on both channel and connectivity parameters. The closing time constant of the ChR2 channel, *τ_OFF_*, governed the duration of the response. For longer *τ_OFF_*, as in Witt *et al*. (Witt et al., 2013), the duration of the evoked spiking and LFP responses in our spiking network model extended well beyond the end of the light pulse. This was because the ChR2 channels slowly closed after the light turned off, allowing excitatory currents to continue for an extended period of time. To account for the differences in ChR2 and ChR2(H134R) (Lin et al., 2009) channel kinetics and their temperature-dependence (Feldbauer et al., 2009) we fit an exponential model to temperature-dependent channel closing times and computed the *in vivo* closing time constant for ChR2(H134R) as approximately 3 ms (**Figure 4—Figure Supplement 1**). Setting *τ_OFF_* to 3 ms generated a response that did not have responses that extended well beyond the offset of the light pulse (**Figure 4C-E**).

### Driven coherent neuronal dynamics depend on timing of excitatory and inhibitory responses

The spiking model allows us to study the dynamics of intracellular currents during stimulation. We estimated the mean driven excitatory and inhibitory currents in the neurons in the model by taking the stimulation pulse triggered average of AMPA-mediated currents, 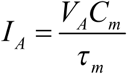 and GABA-mediated currents, 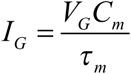, respectively (**Figure 5A**). As the stimulation pulse duration increases, the driven excitatory and inhibitory currents persist for a longer period of time (**Figure 5A**). It is important to note that the E-I windows were not explicitly built into the model, but rather emerged as responses to simulated optogenetic stimulation. As we increased stimulation pulse duration, the relative timing of both the onset and offset of the excitatory and inhibitory currents remained the same, with excitatory currents leading inhibitory currents (**Figures 5B,C**). Thus, the frequency content of the driven coherent neuronal dynamics depends on the duration of the E and I responses.

**Figure 5.**
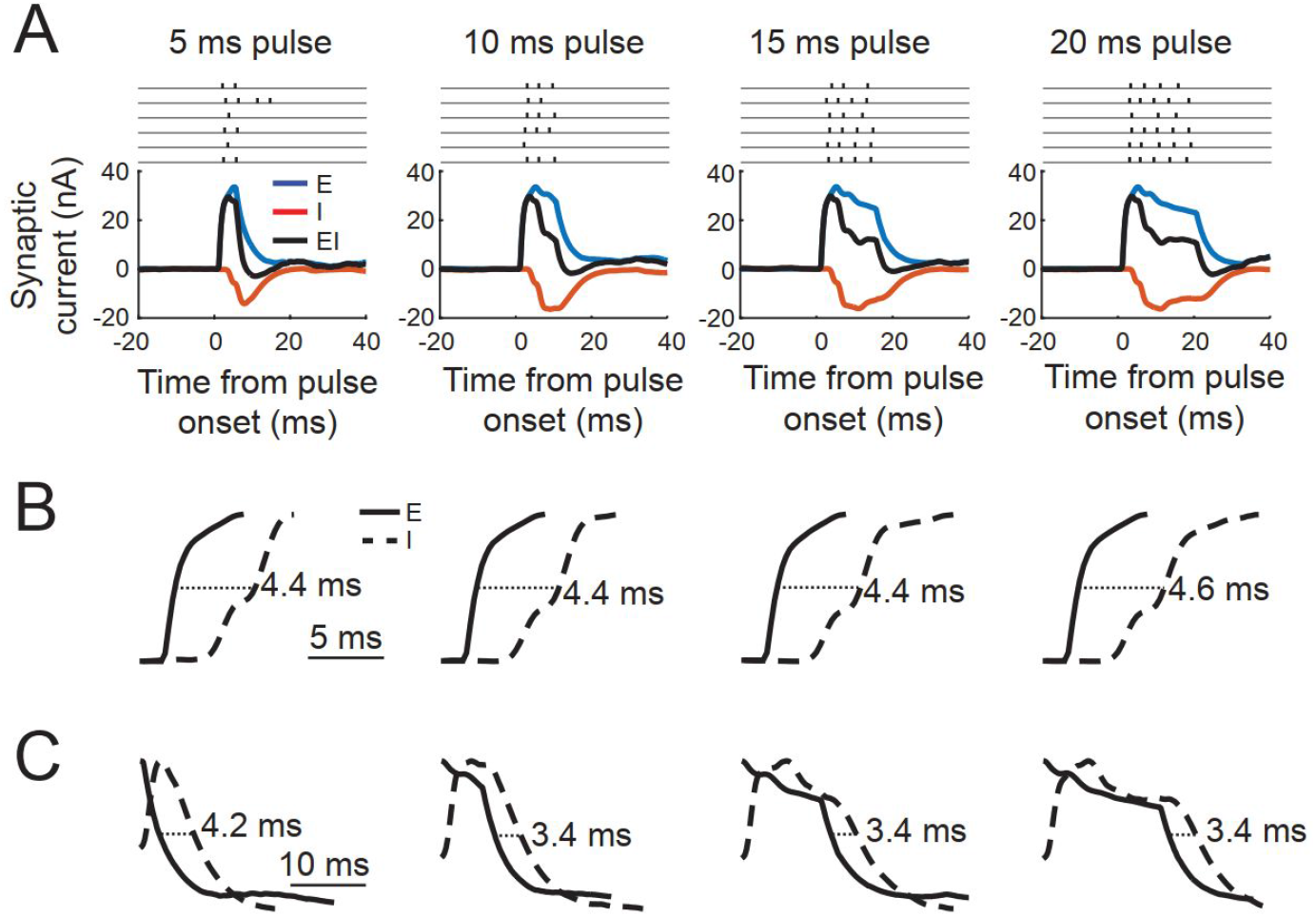
The dynamics of E and I responses varies with pulse duration. **(A)** Stimulation pulse-triggered excitatory (blue), inhibitory (red) and combined E-I responses in neurons in the spiking network model in response to 5, 10, 15 and 20 ms duration simulation pulses. Examples of single pulse evoked spiking are shown above. **(B)** Onset of normalized excitatory (solid) and inhibitory (dashed) responses for 5,10,15 and 20 ms pulse durations. Onset delay is the difference between the time to half maximal amplitude for excitatory and inhibitory responses. **(C)** Offset for normalized excitatory (solid) and inhibitory (dashed) responses for different pulse durations. Offset delay is the difference between the time to half maximal amplitude from the peak of the onset response for excitatory and inhibitory responses.

Previous work has suggested that balanced inhibition is responsible for the temporal precision of spiking in response to sensory stimulation (Wehr and Zador, 2003). In our spiking network model, E-I balance arises from correlations between excitatory synaptic currents and inhibitory synaptic currents during both spontaneous activity and optogenetic stimulation and is thus an intrinsic property of the circuit (**Figures 6A-C**). Consistent with previous experimental results, we find that inhibition lags excitation in the spiking model. Interestingly, the lag decreases during optogenetic stimulation (2.6 ms during spontaneous activity and 1.8 ms during stimulation) (**Figure 6B**) and coherence between E and I activity increases (**Figure 6D**) so that the inhibitory activity more closely tracks the excitatory activity.

**Figure 6:**
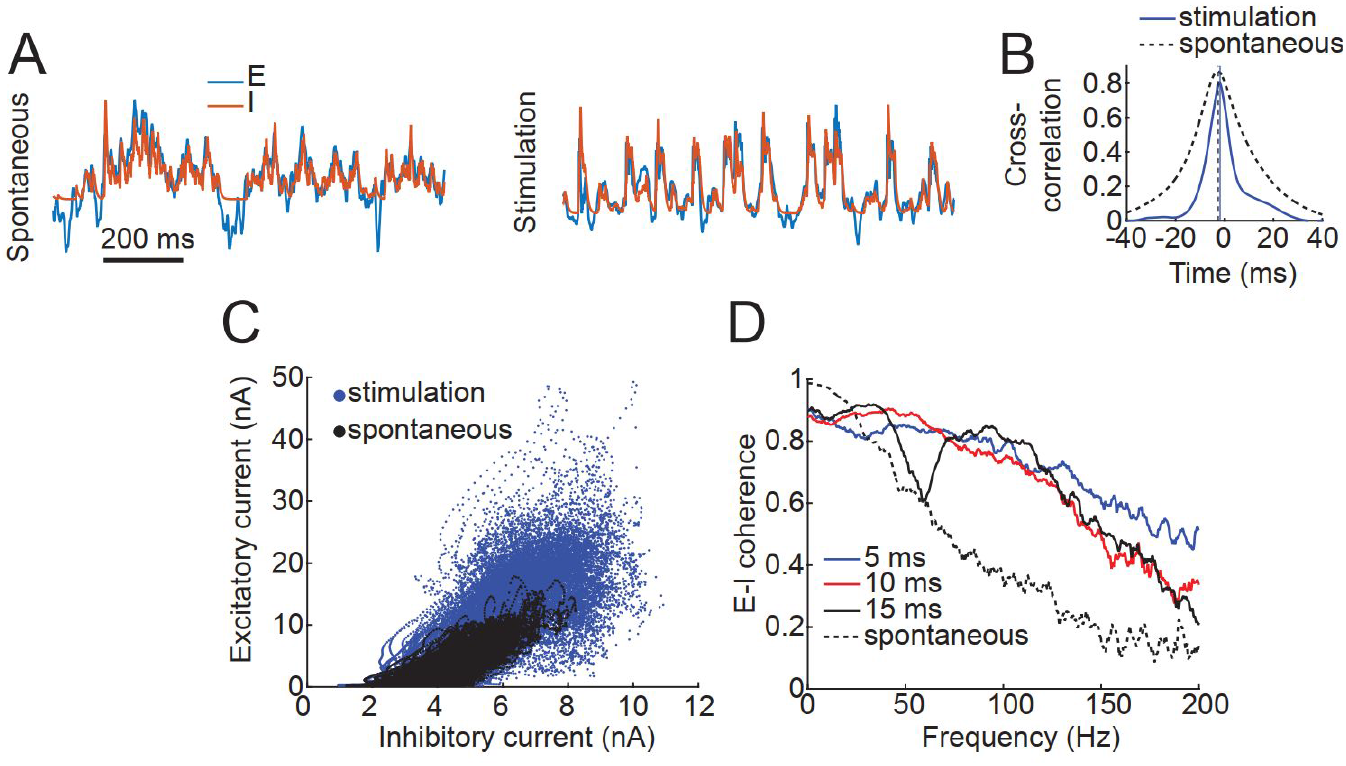
Balanced excitatory and inhibitory responses. **(A)** Example excitatory (blue) and inhibitory (red) synaptic currents in the spiking network model during spontaneous activity, (left), and stimulation,(right). Currents are normalized to show overlap. **(B)** Cross-correlation between E and I synaptic currents during spontaneous activity and the 5 ms stimulation condition. Dotted lines show time lag for maximal cross correlation. **(C)** Instantaneous excitatory and synaptic currents for the 5 ms pulse width condition measured in 0.2 ms intervals. Spontaneous: ρ = 0.82, p = 0; Stimulation: ρ = 0.73, p = 0. **(D)** Coherence between excitatory and inhibitory currents during spontaneous activity (dotted line) and stimulation using 5 ms, 10 ms and 15 ms pulse durations.

The spiking model shows that optogenetic stimulation drives excitatory postsynaptic currents in neurons, which increases their probability of spiking. However, due to the E-I balance inherent in the local network, the excitatory currents are followed closely in time by a balanced amount of inhibitory postsynaptic currents, which suppresses the increased tendency to fire. Therefore, E-I balance, which increases during stimulation, may be responsible for the temporal precision of the driven spiking and thus may be a mechanism for stimulation-driven coherent neuronal dynamics.

## Discussion

Here, we investigate the mechanisms by which optogenetic stimulation of non-human primate (NHP) cortical neurons evokes coherent neuronal dynamics. We show that optogenetically-driven neuronal dynamics arise from how extrinsic inputs recruit excitatory and inhibitory responses that follow intrinsic circuit properties, leading to an effective temporal window for spiking – the E-I window. We develop a phenomenological model to predict how the frequency of optogenetic-stimulation-evoked coherence varies parametrically with stimulation parameters according to the timing of excitatory and inhibitory currents generated by each stimulation pulse. Experimental results from optogenetically-stimulating sites in the posterior parietal and frontal cortex confirmed the model predictions. Furthermore, we developed a biophysical spiking network model of the empirical coherent neuronal dynamics and observed emergent stimulation-driven E-I windows. The biophysical spiking model showed that the E-I windows that shape the extrinsic coherence are an emergent property of the interactions between populations of neurons and indicating that coherence under normal circumstances can also arise from intrinsic E-I balance. Our results contribute to the growing literature on neural coherence by connecting driven dynamics with work on E-I balance.

### E-I window mechanisms of driven coherence

Neurons fire spikes more often during periods of excitatory and inhibitory imbalance, called integration windows. Synchronous excitatory synaptic inputs depolarize cellular membrane potentials and drive spiking activity. Inhibition dominates cortical responses to stimulation (Haider et al., 2013) and regulates the temporal properties of integration windows through temporally lagged responses which balance excitatory inputs and repolarize membrane potentials (Gabernet et al., 2005). Spontaneous excitatory and inhibitory activity are correlated, with inhibition lagging excitation, suggesting that E-I balance may arise from the properties of the neural circuit (Okun and Lampl, 2008). Therefore, both timE-Iagged, correlated excitatory and inhibitory responses may play a key role in spike timing precision in response to optogenetic stimulation, thus generating coherent neuronal dynamics.

The E-I window model describes how excitatory and inhibitory activity affect spike timing and driven coherent neuronal dynamics. To gain further mechanistic insight into the consequences of imbalances in excitatory and inhibitory activity, we developed a biophysical spiking network model. Experimental agreement with the model predictions reinforces the idea that driven coherent neuronal dynamics depends on a mechanism of spike timing precision involving a temporary increase in excitatory activity that leads to an increase in spiking. This is then followed by an increase in inhibitory activity that returns the probability of spiking to baseline. Pulse-duration dependent E-I windows were also an emergent property of a spiking network model of driven coherent neuronal dynamics. Together, these results suggest that the extrinsic generation of E-I windows precisely times spikes and generates coherent neural activity.

The E-I window model also offers a more general mechanism of how spike-field coherence arises intrinsically, in addition to being driven by extrinsic inputs. Several lines of work point to the role of E-I windows in SFC observed under normal circumstances. Sensory stimulation drives E-I responses which govern spike timing (Wehr and Zador, 2003). Since LFP activity reflects temporally structured fluctuations of local excitation and inhibition (Buzsáki and Wang, 2012; Csicsvari et al., 2003; Hasenstaub et al., 2005) LFP activity measures summed local postsynaptic potentials (Buzsáki and Wang, 2012; Pesaran et al., 2018), Thus the intrinsically generated E-I windows are reflected in LFP activity which we demonstrate across multiple regions of the posterior parietal and frontal cortices. This indicates that the E-I window may be a general mechanism for observed coherent neuronal dynamics.

### Relation to cell-type specific manipulations

Previous work suggests that controlling activity in a single cell-type is necessary for driving coherence (Adesnik and Scanziani, 2010; Cardin et al., 2009; Lu et al., 2015; Sohal et al., 2009). However, cell-type-specific stimulation recruits synaptically-mediated responses from other strongly connected neurons in the network (Phillips and Hasenstaub, 2016). The frequency content of coherent neuronal dynamics is shaped by the recruited neuronal ensemble (Kato et al., 2017). This suggests that neuronal coherence evoked by cell-type-specific optogenetic stimulation may also drive an effective E-I window. Our work shows that pan-neuronal optogenetic stimulation generates E-I windows that drive coherent neuronal dynamics. Whether the E-I window model is as effective in predicting the impact of cell-type-specific manipulations that target either E or I cells alone is an open question and will likely depend on the strength of recurrent connectivity in the network.

### Non-oscillatory dynamics

Many computational models of the mechanisms of coherent neuronal dynamics have focused on simulating periodic, oscillatory activity (Börgers and Kopell, 2003; Brunel and Wang, 2003; Keeley et al., 2019; Wilson and Cowan, 1972). However, empirically observed neuronal coherence is episodic (Burns et al., 2011, 2010; Feingold et al., 2015; Rule et al., 2017; Sherman et al., 2016; Xing et al., 2012), with phase and frequency varying over time, and can be narrow-band or broad-band. The computational models we present expand on these prior models to describe manipulations of empirical coherent neural activity patterns which are not necessarily oscillatory or narrow-band and are episodic. Therefore, our models suggest that the interaction between extrinsic inputs and intrinsic circuit properties can drive episodic dynamics in addition to oscillatory dynamics shown in previous models.

### Poisson stimulation for studying dynamics

Continuous and periodic pulse optogenetic stimulation sequences have previously been used generate beta and gamma activity and study their effects on behavior (Adesnik and Scanziani, 2010; Cardin et al., 2009; Ni et al., 2016; Sohal et al., 2009). These forms of stimulation implicitly control the E-I window using the properties of the stimulation pulse and stimulation sequence. However, using continuous constant and pulsatile periodic stimulation drives responses at only particular frequencies (Adesnik and Scanziani, 2010; Cardin et al., 2009; Lu et al., 2015), likely related to properties of the local network (Buzsáki and Wang, 2012) and thus may vary across brain regions and subjects (Lu et al., 2015; Ni et al., 2016). The impact of Poisson stimulation depends on the temporal properties of the stimulation-generated E-I windows rather than the pulse sequence parameters, which we show is shared across multiple cortical regions. Thus an advantage of Poisson stimulation is that it can generate coherent activity across a broader range of behaviorally-relevant frequencies by varying the E-I windows using different stimulation pulse durations in a manner that is general across stimulation sites.

### Potential bias in estimates of coherent neuronal dynamics from LFP activity

Extracellular recordings of LFP activity and spiking may be biased toward activity in pyramidal neurons. LFP activity is most sensitive to postsynaptic potentials in neurons with separated current sources that generate an open field geometry (Pesaran et al., 2018). Thus, contributions from neurons with extended dendritic architectures, such as pyramidal neurons, are more heavily weighted by LFP activity. Additionally, pyramidal neurons typically have larger cell bodies than other cortical neurons and thus have more easily measurable action potentials. Therefore, our measurement of both spiking activity and LFP, and thus SFC, may be biased toward contributions from pyramidal neurons. In addition to LFP activity, alternative decompositions of neural data, such as projections of population activity onto lower dimensional subspaces, may be used to estimate coherence. Such estimates may be more effective at identifying spiking that is coherent with pre- and postsynaptic activity in neuronal ensembles containing both open and closed field neurons.

In conclusion, we systematically modeled neuronal responses to optogenetic stimulation in NHP cortex. Our empirical results highlight the role of generating specific E-I windows by selecting stimulation parameters—pulse duration—to selectively perturb behaviorally-relevant coherent neuronal activity. Our modeling results show that E-I windows that shape the extrinsic coherence are an emergent property of the interactions between populations of neurons and indicate that coherence under normal circumstances can also arise from intrinsic E-I balance. Therefore, optogenetic control of neuronal activity can be useful for understanding mechanisms of coherent neuronal dynamics and how they give rise to complex, flexible behaviors.

## Methods

### Animals

Recordings were taken from two male rhesus macaques (*Macaca mulatta*) (Monkey J: 12 years, 6.5 kg; Monkey H: 11 years, 10.5 kg) and one male cynomolgus macaque (*Macaca fascicularis*) (Monkey B: 4 years, 5 kg). Monkeys J and H had been part of previous studies using the contralateral intraparietal sulcus and the ipsilateral frontal cortex. Monkey B had not been part of previous experiments. All animals were in normal health and were group housed with other conspecifics at the facility. All surgical and animal care procedures were approved by the New York University Animal Care and Use Committee and were performed in accordance with the National Institute of Health guidelines for care and use of laboratory animals.

### Injection surgery

We performed injections of optogenetic viral vectors 6-25 weeks prior to recording. Subjects were maintained at a constant depth of anesthesia with isofluorane (1-2%). Monkey J and H were injected in existing craniotomies. In Monkey B, after mobilizing the scalp and other soft tissue, we made craniotomies over motor cortex to provide access for injections. Injections were made through 26 s gauge stainless steel cannulas (Hamilton) that were inserted through a thinned dura. For each injection, one microliter of AAV5-hSyn-ChR2(h134R)-EYFP was injected at a rate of 0.05 μl/min. We made injections at 2-3 depths spaced 500-750 μm across 2-4 cortical sites per animal. We injected at multiple sites simultaneously to increase the total number of injections sites and thus increase the probability of having successful transduction at one or more sites. We waited 20 minutes between each injection to allow for the virus to diffuse through the tissue. After injections, chambers were reclosed with the chamber cap and periodically opened for cleaning and maintenance while the viral vector was expressing.

### Optical Stimulation

We illuminated the injection site with a 473 nm Laser (Shanghai Sanctity Laser) connected to a multi-mode fiber optic cable (200 μm core, BFL-200, Thor Labs). One end of the fiber optic cable was terminated with an FC/PC connecter and connected to a collimator (PAFA-X-4-A, Thor Labs) for focusing light from the laser. At the other end of the cable we removed a small amount of buffer with a microstripper (T12S21, Thor Labs) and cleaved it with a diamond knife (S90R, Thorlabs) to ensure a flat edge for minimal power-loss.

The fiber optic was placed on top of the thinned dura. The cable was held in the microdrive and placed in the same guide tube as the recording electrode. We did not insert the fiber into the brain in order to minimize damage to the neural tissue. Light power at the tip varied from 16-20.5 mW. (510-653 mW/mm^2^), as measured prior to each experiment using a power meter (PM100D, Thor Labs). Evoked potentials were visually inspected to ensure that there were no signs of artifacts from the Becquerel effect, including rapid onset or offset of neural responses locked to the timing of the light (Cardin et al., 2010). Light based artifacts typically have a capacitive charging and discharging profile, which has a rapid onset and offset tightly coupled to the onset and offset of the device. We ensured that the onset of the response and the inflection point were not aligned with the times when the light was turned on and off. The neural response is slower than the duration of the light pulse and thus continues to increase for a brief period after the light has been turned off. The onset is delayed because the ChR2 channel dynamics are slower than the chemical response and thus the membrane potential does not start to change immediately. The initial response recruits network activity, which has a slow time course. This is particularly evident for short light pulses (<=5 ms). When there is no artifact, the response inflection point occurs after the light is turned off. This is less clear for longer pulse widths (>15 ms) because adaptation and other properties of channel kinetics can lead to a decrease in the response prior to the end of stimulation.

Light pulse patterns were generated by proprietary software running in LabVIEW (National Instruments) and converted to analog signals using a DAQ card (USB-6251, National Instruments). Stimulation trials consisted of 1 s long bursts of square wave pulses. For each block of trials, we controlled three stimulation parameters: pulse width, pulse rate and pulse distribution. We varied pulse widths from 5-20 ms, pulse rates from 5-20 Hz and used both Poisson and periodic pulse distributions.

In rodents, both measurement and simulation have proposed that ~1% of light power density remains after light has passed through approximately 1 mm of brain tissue (Aravanis et al., 2007; Huber et al., 2008; Yizhar et al., 2011). However, primate brain tissue may have slightly different optical properties, thus altering the light transmission. In addition, we stimulated transdurally which may have additional effects on the light. However, we were able to record purportedly directly driven spiking (low-latency, low-jitter) as far as 1.5mm below the cortical surface (not shown), suggesting that we were activating neurons into layer 3.

We tested each recording site in pseudo-randomly interleaved blocks of recordings. Each block consisted of 30-60 repetitions of 1 s long sequences of square wave pulses (bursts), with 1-3 s of no stimulation between each stimulation burst (**Fig 2C**). We held other stimulation parameters (pulse rate, duration and intensity) constant within each block.

Poisson and periodic stimulation pulse trains were generated *in silico* using the same pulse width and rate parameters. Simulations consisted of 1000 trials for each combination of parameters.

### Electrophysiology

We recorded neural signals from awake, alert macaques while they were seated, head-fixed in a primate chair (Rogue Research) in a dark, quiet room. Neural signals were recorded on glass-coated tungsten electrodes (Alpha-Omega, impedance: 0.7-0.8 MO at 1 kHz) held in a FlexMT microdrive (Alpha Omega). Neural signals were acquired at 30 kHz on an NSpike DAQ system and recorded using proprietary data acquisition software. Neural recordings were referenced to a metal wire placed in contact with the dura and away from the recording site. We recorded driven LFP activity and multi-unit spiking in premotor cortex in monkey B (pre-motor cortex) and posterior parietal cortex in Monkeys J and H.

### Fitting E-I Windows

E-I windows were fit to the form

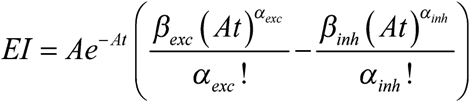

which has an excitatory component

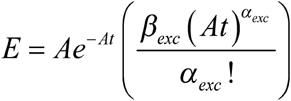

and an inhibitory component

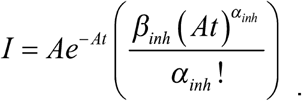

For the pure simulations, *A, β_exc_* and *β_inh_* were set to 1. To fit empirical E-I windows, all parameters were fit using a grid search to minimize the squared error between the modelled window and the empirical window.

### E-I Window Model

We model neuronal spiking according to a conditional intensity function, *λ*(*t*), which represents the instantaneous firing rate, such that the probability of a spike occurring at time *t* is given by *λ*(*t*)Δ*t* (Brown et al., 2003). In the absence of optogenetic stimulation, we model background spiking activity as a Poisson process with a constant rate, *λ*_0_ (**Figure 1A**). We model background LFP activity as a process with increased power at low frequencies–“brown noise”. For simplicity, we further assume that background LFP activity is uncorrelated with spiking activity and does not affect the value of *λ*(*t*).

To simulate LFPs using an excitation window model, we generated 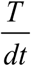 samples of Brownian noise as background activity, with *T* =1 s and *dt* =1 ms. We estimated an E-I window for each recording site by fitting the stimulation pulse evoked LFP response to the following model:

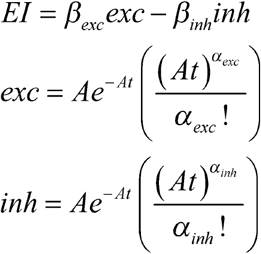

We simulated the driven LFP by convolving the site-specific E-I window with a sequence of delta pulses determined by the stimulation parameters. The driven LFP was and added to the background activity to generate the total simulated LFP activity.

In the model, separate E and I responses to stimulation pulses are summed to create the E-I window. For illustrative purposes, we show the E response preceding the I response but this can be reversed so that I precedes E (Cruikshank et al., 2010; Haider et al., 2013). The E-I window is then convolved with a pulse train to model the combined E-I response to stimulation. To simulate spiking responses, the positive- and negative-rectified responses to the pulse train are passed through a non-linear transformation and summed to generate the time varying firing rate, *λ*(*t*). Individual action potentials are then randomly generated at each time point from a binomial random variable with *p*(*spike at time t*) = *λ*(*t*) Δ*t*.

We modelled spiking activity that was coherent with the LFP using a Poisson point process with a conditional intensity function. The conditional intensity function models stochastic spiking governed by a time-varying instantaneous firing rate modelled as

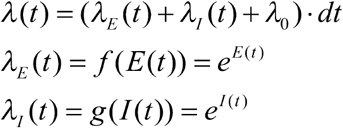

where *λ*_0_ is the spontaneous firing rate, and *dt* is the sampling rate. *E*(*t*) and *I*(*t*) are the time courses of optogenetically driven E and I response, respectively, that generate the fluctuations in the conditional intensity function, *λ*(*t*). *E*(*t*) and *I*(*t*) is determined by convolving the E-I window with a Poisson process generated from the stimulus sequence parameters, then positively or negatively rectifying the E-I response to get the separate E and I responses. After a spike we used a fixed refractory period by forcing *λ*(*t*) = 0 for 3ms.

Simulations of the excitation window model were run using custom scripts in MATLAB using a 1 ms time step. For each trial, we simulated 1 s of spontaneous activity and 1 s of stimulation. We simulated 1000 trials for set of stimulus parameters.

### Spiking network model

The spiking network model consists of a network of 4000 excitatory and 1000 inhibitory leaky integrate-and-fire neurons, each of whose membrane potential, *V*, is described by the equation:

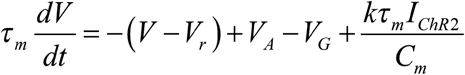

Neurons are randomly connected to each other with a connection probability *P*_*e/i,e/i*_. In addition, the neurons receive connections from external inputs (described below), which represent synaptic input from spontaneous activity in other areas of the brain. The synaptic potentials, *I_A_* and *I_G_*, taken from the Mazzoni model, represent the sum of synaptic responses of each neuron induced by individual pre-synaptic spikes from external inputs and other excitatory (pyramidal, AMPA) and inhibitory (interneurons, GABA) neurons in the model.

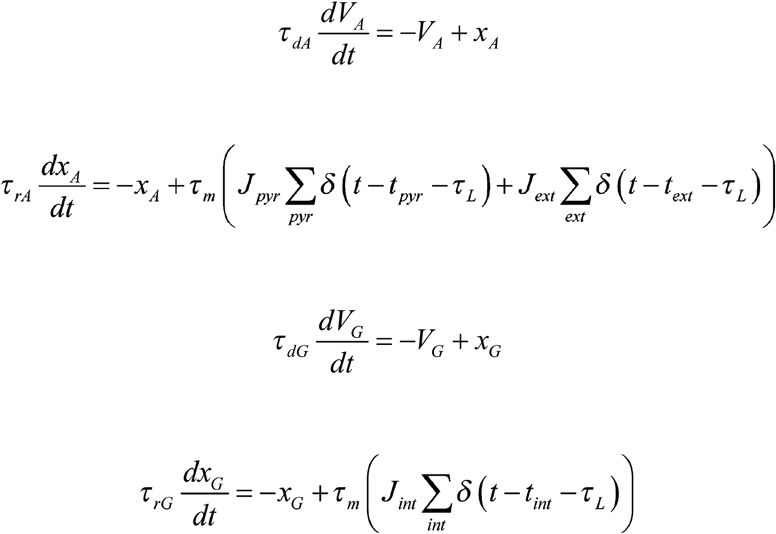

In the model, *t_pyr/int/ext_* is the time of a spike on a presynaptic neuron. The synaptic currents generated by pre-synaptic spikes have latency *τ_L_* and have dynamics governed by their respective exponential rise times (*τ_rA_,τ_rG_*) and decay times (*τ_dA_,τ_dG_*). The efficacy of each connection type is given by *J_pyr/int/ext_* and is uniform for each neuron type. **Table 1** shows the parameters for the Mazzoni part of the model.

**Table 1:**
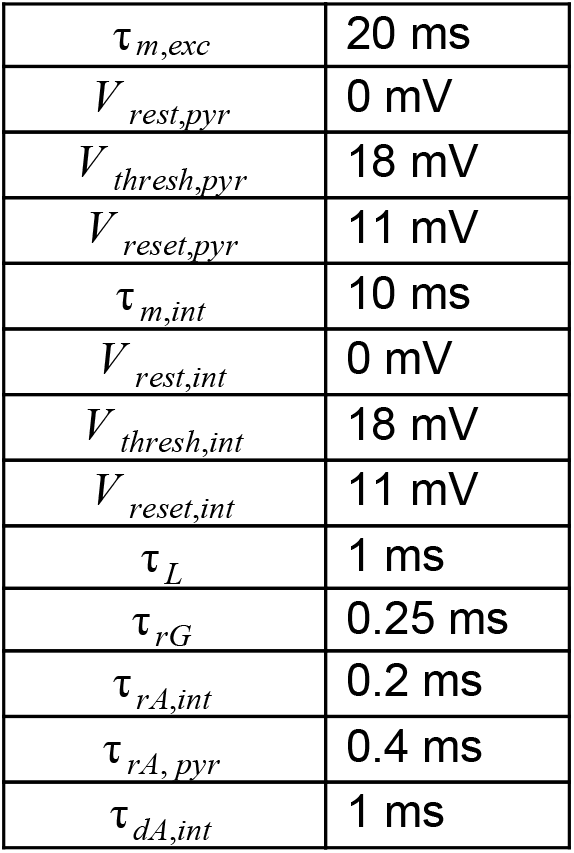
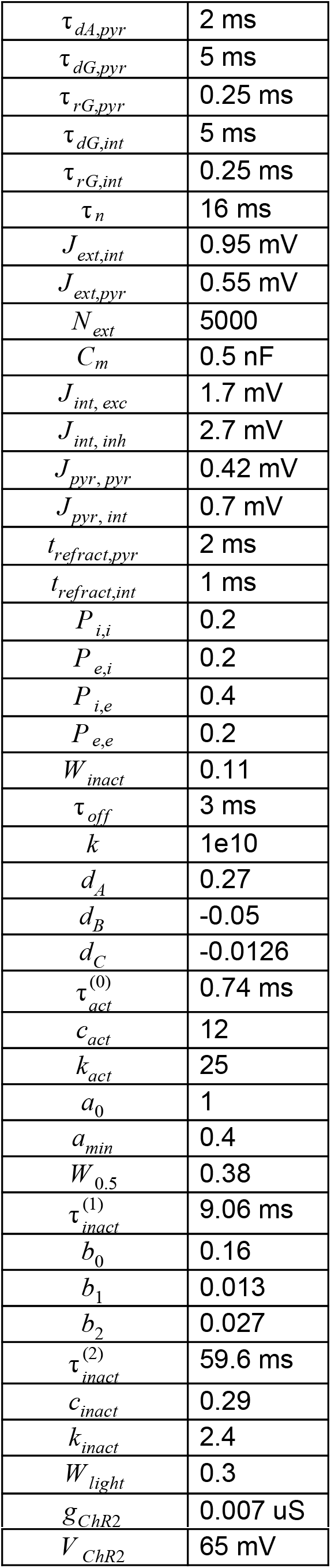
Spiking network model parameters.

#### External Inputs

We modeled spontaneous external activity as inputs generated from excitatory neurons having random Poisson spike trains with a time-varying rate, *r_ext_*(*t*), that is the same for all neurons (Mazzoni et al., 2008). The spike trains are generated from a Poisson process with a rate governed by an Ornstein-Uhlenbeck process:

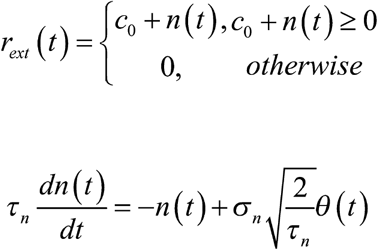

where *θ*(*t*) is Gaussian white noise, *c*_0_ is a non-time-varying mean firing rate and *r_ext_*(*t*) is half-wave rectified to ensure that all firing rates are non-negative.

#### Channelrhodopsin photocurrent

We modeled the channelrhodopsin photocurrent in the transduced neurons according to a model from Witt *et al*. (Witt et al., 2013).

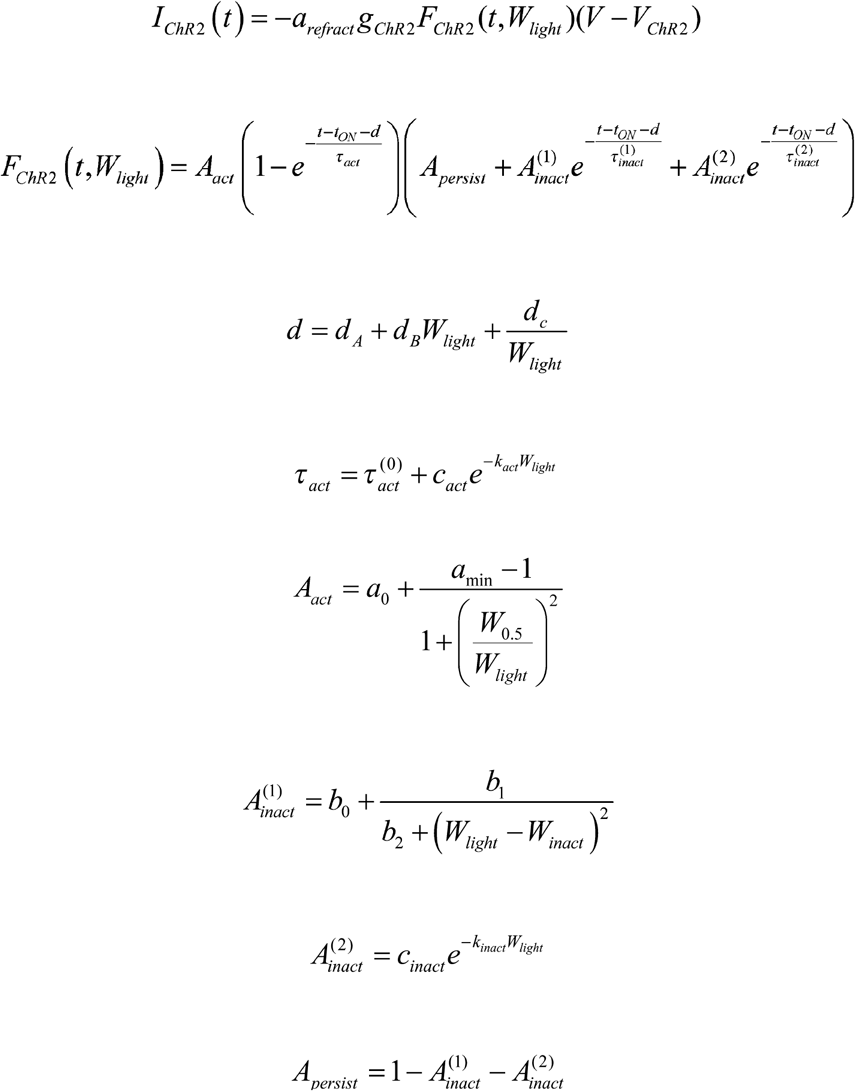

The photocurrent, *I_ChR2_*, is defined as the product of the conductance, *g_ChR2_ F_ChR2_*(*t, W_light_*), and the driving force, *V–V_ChR2_*, from the ChR2 channels at any time, *t*. When light stimulation is on, the conductance varies over time as described by the function *F_chR2_*. When the light is off, the conductance returns to baseline following a single exponential decay with a time constant of *τ_OFF_*. The light induced conductance change is described as the product of activation and inactivation functions which depend on the time from light onset *t–t_ON_*, a light intensity dependent delay, *d*, and their respective time constants, *τ_act_*, 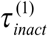 and 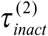. The parameters for equations 8-15 have been previously fitted to photoconductance measured *in vitro* human embryonic kidney cells transduced with AAV1/2-CAG-ChR2-YFP (Witt et al., 2013). **Table 1** shows the parameters for the ChR2 current part of the model. Each neuron of each type was randomly assigned to have ChR2 channels according to the transduction probability for that cell type. We used a viral vector containing the pan-neuronal hSyn promoter. Therefore, we set equal transduction probabilities equal to 1% for both excitatory neurons and inhibitory neurons.(Diester et al., 2011).

Previous studies have shown that transduced neurons do not have robust spiking responses to individual light pulses at higher stimulation frequencies (Lin et al., 2009), which is likely due to the slower kinetics of ChR2 channels. This response fidelity drop-off tends to start around 20 Hz periodic stimulation and reaches near zero around 80-100 Hz stimulation (Cardin et al., 2009). Therefore, light pulses that occurred less than 50 ms after the previous pulse activated ChR2 channels with a probability that depended linearly such that

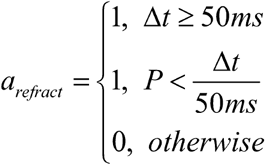

where *P* ∈ *unif*(0,1) and is chosen individually for each close pulse occurrence and for each model neuron.

### Pre-processing in vivo data

Broad-band neural activity was split into multi-unit signals and local field potentials (LFP) by high-pass and low-pass filtering, respectively, at 300 Hz. LFP signals were down-sampled to 1 kHz. Multi-unit activity (MUA) was computed by applying a 3.5 standard deviation threshold to the multi-unit signals to identify putative spikes. All waveforms that exceeded this peak were labelled as multi-unit action potentials.

### Pre-processing in silico data

LFPs measure currents generated by synaptic activity within a volume near the electrode (Logothetis, 2003). Therefore, we followed Mazzoni *et al*. (Mazzoni et al., 2008) and modeled the LFP as a sum of synaptic currents. Since interneurons typically have dendrites that are organized in a symmetrical, closed-field configuration, we assumed their contribution to the LFP would be minimal in comparison with synaptic currents on the typically dipole-like dendrites of pyramidal neurons (Buzsáki and Wang, 2012). Therefore, only synaptic currents in pyramidal neurons contributed to our measure of the model LFP. Adding together the absolute values of synaptic currents has been shown to accurately reproduce the power spectrum and information content of LFP recorded in macaque visual cortex (Mazzoni et al., 2008). Simulated LFPs were also low-pass filtered at 300 Hz and down-sampled to 1 kHz. Simulated multiunit activity consisted of all the threshold crossings in the leaky integrate-and-fire model, as recorded by the Brian simulator. The *in vivo* and *in silico* data were pre-processed, were treated the same after pre-processing.

### Data Analysis

We computed multi-unit peri-stimulus time histograms (PSTH) by aligning MUA to the onset time of light pulses. Individual spikes were collected in 1 ms bins and the corresponding histogram was smoothed by convolution with a Gaussian function with a standard deviation of 1 ms. In order to avoid corruption from nearby pulses, evoked potentials were computed by averaging LFPs aligned to the onset time of light pulses only for pulses that occurred at the end of a pulse train and did not have a pulse preceding it by less than 50 ms.

Spectral analysis was performed using multi-taper methods (Mitra and Pesaran, 1999). The accuracy of the spectral quantities we used does not depend on the center frequency of the estimate for all frequencies above the minimum resolvable frequency, given by the frequency smoothing in Hz (Jarvis and Mitra, 2001). For each recording, power spectra were computed using a 1 s analysis window and ± 1.5 Hz frequency smoothing. We computed power spectra for the LFP responses during entire 1 s burst (stimulation epoch) and for 1 s prior to the burst (spontaneous epoch). Power spectra were computed for neural signals for the spontaneous and stimulation epochs. Recording sites with power spectra that did not exhibit a significant difference were not included in analysis. The exclusion criteria for recording sessions did not depend on the frequency content of the driven LFP response—the desired effect—but rather only that there was any driven LFP response at all. Sites were excluded if there was no change in the LFP power between spontaneous and stimulation periods for the 10ms pulse width condition, which was likely due to the electrode being placed on the periphery of light cone and thus failing to record driven neural activity. We explicitly confirm this in our experiments by moving the electrode and observing driven responses. We computed the spectrogram for each recording using a ± 250 ms analysis window, stepped 50 ms between windows, with ± 2.5 Hz frequency smoothing aligned to the start of the burst. We computed the spike-field coherence for each recording using a 1 s analysis window with ± 3 Hz frequency smoothing averaged across at least 30 repetitions.

## DATA AND SOFTWARE AVAILABILITY

Software is available in a GitHub repository linked from http://www.pesaranlab.org. Data are available upon reasonable request.

## Acknowledgements

This work was supported, in part, by US National Institutes of Health (NIH) BRAIN Initiative grant R01-NS104923, NIH R01-EY024067, NIH P30-EY013079, NIH Training Grant T32-EY007136, a Scholar Award from the McKnight Endowment Fund for Neuroscience (BP) and the Alfred P Sloan Foundation (BP).

## Author Contributions

Conceptualization: R.A.S., H.L.D., Y.T.W., M.A.H., and B.P.; Methodology: R.A.S., H.L.D., Y.T.W., M.A.H., M.M.F. and B.P.; Software: R.A.S. and B.P.; Formal Analysis: R.A.S. and B.P.; Investigation: R.A.S. and B.P.; Resources: R.A.S., H.L.D., and B.P.; Writing – Original Draft: R.A.S. and B.P.; Writing – Review & Editing: R.A.S., H.L.D., Y.T.W., M.A.H., M.M.F. and B.P.; Visualization: R.A.S. and B.P.; Supervision: H.L.D., Y.T.W., M.A.H., and B.P.; Funding Acquisition: B.P.

---

**Figure 3—figure supplement 1.**
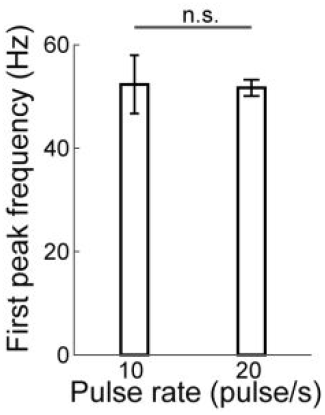
The response dynamics do not depend on pulse rate. Average and SEM of the standard deviation of the first fitted Gaussian for all stimulus sequences with a 10 ms pulse width across all recording sites.

**Figure 4—figure supplement 1.**
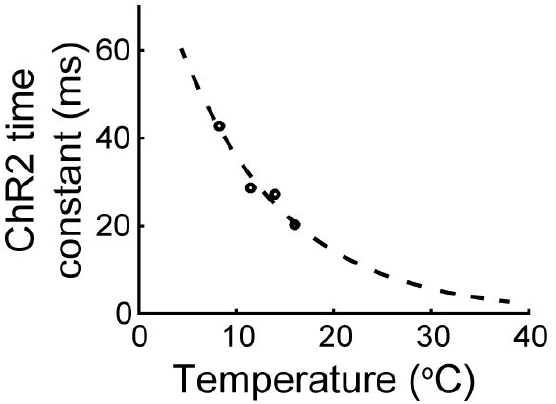
Fitting ChR2 channel kinetics to *in vivo* temperatures. Approximate ChR2 channel kinetics adapted from Feldbauer et al. with an exponential fit and extrapolation to macaque body temp (about 37° F).

